# Expanding N-terminomics Coverage by Extending N-terminal Peptide Length

**DOI:** 10.64898/2026.09.22.753420

**Authors:** Akane Takeshita, Eisuke Kanao, Koshi Imami, Yasushi Ishihama

## Abstract

In bottom-up proteomics, protease specificity determines peptide length and LC/MS/MS detectability; in N-terminomics, this is critical because each protein N-terminal peptide is unique and cannot be substituted by other peptides. CHromatographic AMplification of Protein N-terminal peptides (CHAMP-N) combines LysargiNase digestion with StageTip-based strong cation exchange chromatography (SCX), removing internal peptides by exploiting their newly generated N-terminal Lys/Arg residues and thereby selectively isolating protein N-terminal peptides. However, LysargiNase cleavage near the protein N-terminus can generate excessively short peptides. Here, we developed two complementary CHAMP-N approaches that modurate N-terminal peptide length while retaining the charge-based SCX separation principle. LysN digestion avoided cleavage at N-terminal-proximal Arg residues, whereas D3-acetylation blocked Lys cleavage by LysargiNase to produce ArgN-like specificity. Both approaches extended otherwise short N-terminal peptides while preserving N-terminal basic residues on internal peptides. Conventional LysargiNase, LysN, and ArgN-like CHAMP-N generated complementary sets of N-terminal peptides. Integration of the three workflows identified 2,552 protein N-termini, approximately twice the number identified by conventional CHAMP-N alone. The median relative standard deviations of N-terminal peptide intensities were approximately 15% for all three workflows, indicating comparable technical reproducibility. These results demonstrate that coordinating protease cleavage specificity with SCX selectivity reduces sequence-dependent losses and expands N-terminome coverage while preserving the simplicity and reproducibility of CHAMP-N.

## Introduction

Protein N-termini are information-rich regions that reflect protein maturation states and intracellular fates. Initiator methionine removal and Nα-acetylation are cotranslational events that occur in many proteins,^1,2^ and the resulting N-terminal residues and their modification states also contribute to the regulation of protein stability through N-degron pathways.^3,4^ In addition to transcript diversification through selective promoter and transcription start site usage and alternative splicing, noncanonical translation initiation at upstream or downstream AUG codons or near-cognate codons generates proteoforms with extended or truncated N-termini.^5,6^ Proteolytic processing further generates neo-N-termini,^7^ while cleavage of signal peptides and organelle-targeting sequences exposes the N-termini of mature proteins.^8,9^ Through these transcriptional, translational, and post-translational processes, a single gene can give rise to multiple proteoforms with distinct N-termini.^10,11^ Therefore, N-terminal analysis by bottom-up proteomics (N-terminomics), which enables large-scale profiling of N-terminal sequences, is important for understanding proteoform diversity that cannot be captured by gene annotation alone.

Conventional N-terminomics enriches N-terminal peptides either by exploiting differences in amino-group state, charge, or hydrophobicity between protein N-terminal peptides and internal peptides generated by enzymatic digestion or by selectively introducing an affinity tag at the N-terminus. Representative negative-selection methods include COmbined FRActional DIagonal Chromatography (COFRADIC)^12,13^ and Terminal Amine Isotopic Labeling of Substrates (TAILS),^14,15^ and their enrichment principles and derived methods have been summarized in a review.^16^ Positive-selection methods include subtiligase-mediated enzymatic ligation of a biotin-containing peptide to free N-termini^17^ and direct labeling and capture of free N-termini using 2-pyridinecarboxaldehyde.^18^ More recent advances include methods applicable to low-input samples,^19,20^ parallel processing of multiple samples,^21,22^ and an ion mobility-based approach that eliminates the need for prior enrichment.^23^ However, many N-terminomics workflows require multistep procedures involving predigestion, chemical modification, chemoselective or affinity-based capture, and fractionation.

To reduce this operational complexity, we developed CHromatographic AMplification of Protein N-terminal peptides (CHAMP-N), which combines LysargiNase digestion with strong cation exchange chromatography (SCX) separation.^24^ Internal peptides generated by LysargiNase begin with Lys or Arg and are therefore strongly retained by the SCX material, whereas protein N-terminal peptides are retained more weakly. By exploiting this charge difference, CHAMP-N isolates protein N-terminal peptides in a single SCX separation without postdigestion chemical derivatization or affinity capture. We subsequently adapted the original HPLC-based SCX separation to an SCX-StageTip format while retaining comparable identification depth and selectivity.^25^ Because SCX-StageTips allow multiple samples to be processed in parallel, the resulting workflow combines operational simplicity with high throughput and high reproducibility.

Despite advances in enrichment technology, N-terminomics still faces fundamental limitations imposed by the length of the N-terminal peptides produced by protease digestion. While global proteomics allows a single protein to be identified by multiple peptides, in N-terminomics, the N-terminus of each protein is represented by only a single peptide. If the cleavage site by the digestive enzyme is too close to the N-terminus of the protein, the resulting N-terminal peptide becomes extremely short; this not only reduces retention in reverse-phase LC and hinders MS/MS identification but also makes it impossible to unambiguously identify the source protein. Conversely, if the cleavage site is too far from the N-terminus, the resulting long peptides may not ionize or fragment adequately. Consequently, if the length of the corresponding peptide is inappropriate, selective enrichment alone cannot guarantee the detection of the N-terminus. Comparisons of theoretical and experimentally observed tryptic peptide-length distributions have shown that peptide length strongly affects MS-based peptide observability.^26^ In addition, in silico trypsin digestion of the human proteome revealed that if N-terminal peptides are too short, insufficient sequence information is obtained to unambiguously identify the corresponding proteins.^27^

Accordingly, strategies that alter effective digestion specificity through chemical modification have been developed to expand the detectable range of terminal peptides. In TAGS, guanidination of Lys side chains and protein N-termini improves tryptic cleavage at modified Lys residues, thereby expanding N-terminal peptide identification.^28^ SAPT combines ArgC-like digestion induced by protein propionylation with SCX fractionation for protein terminus analysis.^29^ These studies demonstrate that control of digestion specificity is an important determinant of terminome coverage.

CHAMP-N is likewise constrained by the cleavage specificity of LysargiNase. Cleavage N-terminal to a Lys or Arg residue near the protein N-terminus can generate an N-terminal peptide that is too short for LC/MS/MS analysis. Conversely, a distant first Lys/Arg cleavage site can generate an excessively long N-terminal peptide with reduced LC/MS/MS detectability. However, theoretical tryptic peptide-length distributions suggest that excessively long peptides are less frequent than excessively short peptides.^26^ We therefore focused on extending N-terminal peptides that would otherwise be shortened by cleavage at N-terminal-proximal Lys or Arg residues. Although proteases with different cleavage specificities can generate complementary N-terminal peptides and expand N-terminome coverage,^30^ their use in CHAMP-N is restricted because its SCX selectivity depends on internal peptides beginning with Lys or Arg. In this study, we investigated whether N-terminome coverage could be expanded using two complementary digestion strategies, LysN digestion and D3-acetylation followed by LysargiNase digestion (hereafter, ArgN-like digestion), while preserving the SCX selectivity of CHAMP-N.

## Materials and Methods

### Materials

LysargiNase was purchased from Merck Millipore (Burlington, MA), and LysN was purchased from ImmunoPrecise Antibodies (Europe) B.V. (Utrecht, the Netherlands). Empore Cation-SR and SDB-XC extraction disks were purchased from GL Sciences (Tokyo, Japan). UltraPure Tris buffer and fetal bovine serum were purchased from Thermo Fisher Scientific (Waltham, MA). Protease inhibitors, acetic anhydride-d6, and hydroxylamine were purchased from Sigma-Aldrich (St. Louis, MO). Water was purified using an ELGA PURELAB^®^ Quest2 water purification system (ELGA LabWater, High Wycombe, UK). Unless otherwise specified, all other reagents were purchased from FUJIFILM Wako (Osaka, Japan).

### Cell culture

HEK293T cells obtained from the RIKEN BioResource Research Center (RIKEN BRC, Tsukuba, Japan) were cultured in Dulbecco’s modified Eagle’s medium (DMEM) supplemented with 10% (v/v) fetal bovine serum at 37 °C in a humidified atmosphere containing 5% CO_2_. The cells were washed twice with ice-cold phosphate-buffered saline (PBS), harvested using a cell scraper, and collected by centrifugation.

### Protein extraction and digestion

Proteins were extracted using a modified phase-transfer surfactant (PTS) protocol.^31^ HEK293T cell pellets were resuspended in PTS lysis buffer containing a protease inhibitor cocktail, 12 mM sodium deoxycholate (SDC), and 12 mM sodium N-lauroylsarcosinate (SLS) in 100 mM HEPES buffer (pH 8.5). The cell suspension was heated at 95 °C for 5 min and sonicated for 20 min. Protein concentrations were determined using a bicinchoninic acid (BCA) protein assay. Proteins were reduced with 10 mM dithiothreitol for 30 min, then alkylated with 50 mM iodoacetamide at 37 °C for 30 min in the dark. For conventional LysargiNase digestion, the protein samples were diluted 10-fold with 10 mM CaCl2 and incubated overnight at 37 °C with LysargiNase at an enzyme-to-protein ratio of 1:100 (w/w). For LysN digestion, the samples were diluted 5-fold with 50 mM ammonium bicarbonate and incubated for 2 h at 37 °C with

LysN at an enzyme-to-protein ratio of 1:200 (w/w). After digestion, an equal volume of ethyl acetate was added, and the samples were acidified to a final concentration of 0.5% (v/v) TFA. The samples were vortexed for 2 min and centrifuged at 15,800 × g for 2 min to separate the aqueous and organic phases. The aqueous phase was recovered and dried in a vacuum concentrator. The peptides were reconstituted in 0.1% (v/v) TFA and desalted using SDB-XC StageTips.^32,33^

### Chemical acetylation

For ArgN-like digestion, protein extraction, reduction, and alkylation were performed as described above. The pH of each protein sample was adjusted to 8.0 with 1 M HCl, and the protein concentration was adjusted to 1 μg/μL. A 50 µL aliquot containing 50 µg of protein was mixed with 1 µL of 0.5 M acetic anhydride-d_6_ in ACN and incubated on ice for 10 min. The reaction was quenched by adding 100 µL of 100 mM Tris-HCl (pH 9.0) and 20 µL of 2% hydroxylamine. The sample was then diluted to a final volume of 500 µL with 15 mM CaCl_2_ and incubated overnight at 37 °C with LysargiNase at an enzyme-to-protein ratio of 1:100 (w/w). Following digestion, the surfactants were removed by phase transfer, and the resulting peptides were desalted as described above.

### Strong cation exchange chromatography

SCX chromatography was performed using disposable pipette tip-based SCX-StageTips, as described previously.^33^ Two membrane disks were punched from Empore Cation-SR disks using a blunt-ended 16-gauge syringe needle and placed in a P200 pipette tip. The loading solution consisted of 0.5% (v/v) TFA and 30% (v/v) acetonitrile. Before use, the SCX-StageTips were conditioned with 30% (v/v) acetonitrile containing 1 M NaCl and equilibrated with 300 µL of the loading solution. An amount of each desalted digest corresponding to 10 µg of protein, as determined by the BCA assay before digestion, was dried in a SpeedVac concentrator and reconstituted in the loading solution. Each sample was loaded onto an SCX-StageTip, and the flow-through and subsequent eluates were collected as specified for each experiment. Pooled protein digests were used throughout the study, and the SCX-StageTip procedure and all subsequent steps were performed in triplicate unless otherwise stated. The collected fractions were concentrated in a SpeedVac and reconstituted in 4% (v/v) acetonitrile containing 0.5% (v/v) TFA. The samples were analyzed using a nanoLC system coupled to an Orbitrap Exploris 480 mass spectrometer (Thermo Fisher Scientific), as described below.

### Comparison of single-step elution and sequential two-step elution

For the two single-step elution conditions, aliquots of the same digest were loaded onto two separate SCX-StageTips. One tip was eluted with 50 µL of 0.5% (v/v) TFA in 30% (v/v) acetonitrile, whereas the other tip was eluted with 50 µL of 5.0% (v/v) TFA in 30% (v/v) acetonitrile. For each tip, the eluate was collected in the same well as its corresponding flow-through, yielding one combined fraction for each elution condition. The combined fractions obtained under the 0.5% and 5.0% TFA conditions were not pooled and were analyzed separately by nanoLC/MS/MS. For sequential two-step elution, each SCX-StageTip was first eluted with 50 µL of 0.5% (v/v) TFA in 30% (v/v) acetonitrile, and the eluate was collected in the same well as the flow-through to generate Fraction 1. The same tip was subsequently eluted with 50 µL of 5.0% (v/v) TFA in 30% (v/v) acetonitrile, and this second eluate was collected separately as Fraction 2. Thus, Fraction 1 comprised the flow-through and the 0.5% TFA eluate, whereas Fraction 2 comprised only the 5.0% TFA eluate. The two fractions were processed independently and analyzed separately by nanoLC/MS/MS as described above.

### NanoLC/MS/MS analysis

NanoLC/MS/MS analyses were performed using an Orbitrap Exploris 480 mass spectrometer coupled to a nanoLC system comprising a WPS-3000PL RS autosampler and an UltiMate 3000 RSLCnano pump (Thermo Fisher Scientific). Mass spectra were acquired in positive-ion data-dependent acquisition mode.

Unless otherwise stated, nanoLC/MS/MS analyses were performed under the following conditions. The LC mobile phases were solvent A, 0.5% (v/v) acetic acid in water, and solvent B, 0.5% (v/v) acetic acid in 80% (v/v) acetonitrile. Peptides were separated on an in-house-packed needle column (250 mm length × 100 µm i.d.) containing ReproSil-Pur 120 C18-AQ resin (1.9 µm; Dr. Maisch). The column was maintained at 50 °C using a PRSO-V2 column oven (Sonation GmbH), and the flow rate was 400 nL min^−1^. The gradient was as follows: 5% B for 5 min, 5–19% B over 55.3 min, 19–29% B over 21 min, 29–40% B over 8.7 min, and 40– 99% B over 0.1 min, followed by 99% B for 4.9 min. The electrospray voltage was set to 2.4 kV. Full MS scans were acquired over an *m/z* range of 375–1500 at a resolution of 60,000, with a normalized automatic gain control (AGC) target of 300% and an automatic maximum injection time. MS/MS scans were acquired at a resolution of 15,000, with the standard AGC target setting, an automatic maximum injection time, and a fixed first mass of *m/z* 120. Precursor ions were fragmented by higher-energy collisional dissociation using a normalized collision energy of 30%. The dynamic exclusion time was 20 s. The cycle time between master scans was 1 sec.

For the experiments presented in Figure 4, the following extended gradient was used: 5% B for 5 min, 5–19% B over 55.3 min, 19–29% B over 21 min, 29–84% B over 43.5 min, and 84– 99% B over 0.1 min, followed by 99% B for 4.9 min. The fixed first mass for MS/MS scans was set to *m/z* 100. All other acquisition parameters were unchanged.

## Data analysis

The raw MS data files were processed using MaxQuant (version 2.6.5.0)^34^, and the database search was performed with Andromeda^35^ against a human UniProtKB/Swiss-Prot protein sequence database including isoforms (2020.10, 42,373 sequences). The precursor and fragment ion mass tolerances were set to 4.5 and 20 ppm, respectively. The minimum peptide length was 7. The maximum peptide mass was 4600 Da. For conventional LysargiNase-based CHAMP-N samples, LysargiNase specificity was selected, and up to two missed cleavages were permitted. For LysN-based CHAMP-N samples, LysN specificity was selected, and up to two missed cleavages were permitted. For both sample types, acetylation of protein N-termini and methionine oxidation were specified as variable modifications, whereas cysteine carbamidomethylation was specified as a fixed modification. For ArgN-like CHAMP-N samples, LysargiNase specificity was selected, and up to five missed cleavages were permitted. Acetylation of protein N-termini, D3-acetylation of protein N-termini (+45.0294 Da), D3-acetylation of lysine residues (+45.0294 Da), and methionine oxidation were specified as variable modifications. Cysteine carbamidomethylation was specified as a fixed modification. The false discovery rate was controlled at 1% at both the peptide-spectrum match and protein levels. For coverage analyses, each leading razor protein associated with at least one identified protein N-terminal peptide was counted as one detected protein N-terminus.

## Evaluating the specificity of D3-acetylation

To evaluate the specificity of D3-acetylation in ArgN-like CHAMP-N samples, the raw data were additionally analyzed by an exploratory open search followed by a targeted variable-modification search using MSFragger (version 4.3) in FragPipe (version 23.1), together with Philosopher (version 5.1.2) and PTM-Shepherd (version 3.0.2).^36–38^ In the targeted search, the +45.0294 Da mass shift was allowed on Ser, Thr, Tyr, Arg, and His in addition to protein N-termini and Lys residues. PSMs were filtered at a 1% FDR. For the site-level analysis, repeated assignments to the same modified site in the same protein were counted only once, and the distribution of the resulting unique sites among protein N-termini and the six residue types was calculated. Protein N-terminal peptides were classified according to their sequences and positions within the corresponding proteins, irrespective of the exact localization of the +45.0294 Da mass shift. Detailed search parameters and calculation procedures are provided in the **Supporting Information**.

## Results and Discussion

### Complementary digestion strategies for modulating N-terminal peptide length in CHAMP-N

To modulate N-terminal peptide length while preserving the SCX selectivity of CHAMP-N, we designed two complementary digestion strategies (**Figure 1**). In Approach 1, LysN digestion was used to avoid cleavage at Arg residues near the protein N-terminus. Because LysN cleaves exclusively N-terminal to Lys, this strategy allows N-terminal peptides to extend to the first downstream Lys. In Approach 2, ArgN-like digestion was achieved by D3-acetylating free amino groups in proteins before LysargiNase digestion. D3-acetylation of the ε-amino groups of Lys suppressed cleavage N-terminal to Lys, allowing N-terminal peptides to extend beyond Lys residues near the protein N-terminus to the first downstream Arg. In both approaches, the resulting internal peptides began with a basic residue, Lys in Approach 1 and Arg in Approach 2, thereby preserving the charge characteristics required for SCX separation in CHAMP-N.

**Figure 1.**
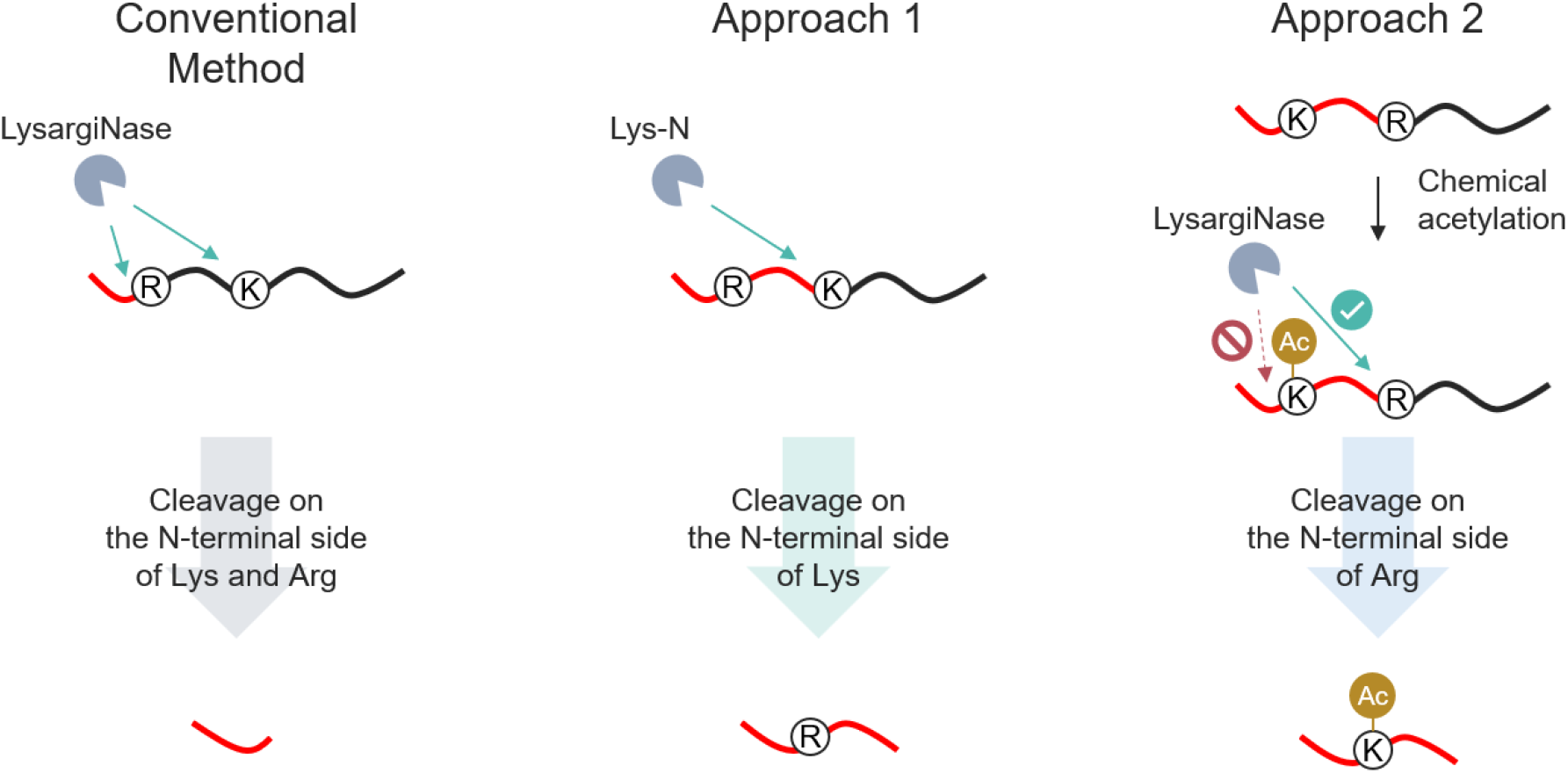
Complementary strategies for extending short N-terminal peptides in CHAMP-N. (Left) In conventional LysargiNase digestion, cleavage occurs N-terminal to both Lys (K) and Arg (R); therefore, the nearest downstream Lys or Arg defines the C-terminal boundary of each protein N-terminal peptide. When this cleavage site lies close to the protein N-terminus, an excessively short N-terminal peptide can be generated. (Middle) In Approach 1, LysN digestion produces cleavage exclusively N-terminal to Lys, thereby bypassing a proximal Arg and extending the N-terminal peptide to the first downstream Lys. (Right) In Approach 2, ArgN-like digestion is achieved by D3-acetylating proteins before LysargiNase digestion. D3-acetylation of the ε-amino groups of Lys suppresses cleavage N-terminal to Lys, allowing the N-terminal peptide to extend beyond a proximal Lys to the first downstream Arg. Both approaches generate internal peptides that begin with a basic residue, Lys in Approach 1 and Arg in Approach 2, thereby preserving the charge characteristics required for SCX separation in CHAMP-N. Red lines indicate protein N-terminal peptides, and Ac denotes D3-acetylation of Lys.

To estimate the theoretical frequency of N-terminal peptides below the database-searchable length range, we performed in silico digestion of canonical human UniProtKB/Swiss-Prot protein sequences (release 2020.10, 20,385 proteins). Cleavage was modeled N-terminal to Lys and Arg, Lys, or Arg for conventional LysargiNase, LysN, or ArgN-like digestion, respectively. No missed cleavages were allowed, and initiator-methionine excision was not considered. The theoretical N-terminal peptide length was defined as the number of residues from the annotated protein N-terminus to the residue immediately preceding the first modeled cleavage site. Because the database searches used the default minimum peptide length of seven residues in MaxQuant (ver. 2.6.5.0), theoretical N-terminal peptides were categorized as either shorter than seven residues or at least seven residues long. This cutoff represents eligibility for the database search rather than an empirical threshold for LC/MS/MS detectability.

Under conventional LysargiNase digestion, 50.6% of the theoretical N-terminal peptides were shorter than seven residues (**Figure S1**). Among the protein N-termini predicted to yield such peptides under conventional LysargiNase digestion, LysN and ArgN-like digestion were predicted to increase the N-terminal peptide length to at least seven residues for 51.8% and 34.5% of these protein N-termini, respectively. Notably, these two groups were mutually exclusive: protein N-termini rescued by LysN digestion had Arg as the first cleavage residue under conventional LysargiNase digestion, whereas those rescued by ArgN-like digestion had Lys. Consequently, the two approaches were predicted to produce N-terminal peptides of at least seven residues for a combined 86.3% of the protein N-termini that yielded peptides shorter than seven residues under conventional LysargiNase digestion. These results indicate that the two approaches theoretically extend complementary subsets of short N-terminal peptides generated by conventional LysargiNase digestion, thereby increasing the proportion that meet the peptide-length criterion for database searching.

### Optimization of SCX Elution Conditions and Expanded Coverage of Protein N-Termini in LysN-Based CHAMP-N

Because LysN does not cleave N-terminal to Arg, protein N-terminal peptides generated by LysN digestion may contain internal Arg residues. Under acidic conditions, such peptides carry additional positive charges due to protonation of the Arg side chains and may therefore be strongly retained by SCX and insufficiently recovered by conventional elution with 0.5% TFA. We therefore fixed the sample-loading conditions at 0.5% TFA and 30% ACN and compared the recovery of N-terminal peptides from LysN digests using elution solutions containing 0.5%, 1.0%, 2.5%, or 5.0% TFA (**Figure S2**). The number of identified protein N-termini increased with increasing TFA concentration. In contrast, N-terminal peptide selectivity, defined as the summed MS intensity of N-terminal peptides divided by the summed MS intensity of all identified peptides, decreased. Thus, elution with 5.0% TFA expanded the recovery of strongly retained N-terminal peptides but also increased the coelution of internal peptides. Notably, increasing the TFA concentration from 0.5% to 5.0% in LysargiNase-based CHAMP-N did not appreciably increase the number of identified protein N-termini, suggesting that stronger elution alone was insufficient to expand coverage under the conventional digestion conditions. Based on these results, we selected 0.5% TFA, which provided high selectivity, and 5.0% TFA, which yielded the largest number of identified protein N-termini, for further evaluation of elution formats.

We next compared single-step elution, in which either 0.5% or 5.0% TFA was applied once, with sequential two-step elution, in which 0.5% and 5.0% TFA were applied sequentially to the same SCX-StageTip (**Figure 2A**). For single-step elution, the same LysN digest was divided into two aliquots and loaded onto separate SCX-StageTips. One StageTip was eluted with 0.5% TFA and the other with 5.0% TFA. For each condition, the flow-through and corresponding eluate were combined into a single fraction and analyzed in one nanoLC/MS/MS run. For sequential two-step elution, a single sample aliquot was loaded onto an SCX-StageTip. The flow-through and 0.5% TFA eluate were collected as Fraction 1, after which the same StageTip was eluted with 5.0% TFA to obtain Fraction 2. The two fractions were analyzed separately by nanoLC/MS/MS, and the identification results were combined. Sequential two-step elution recovered, from a single sample aliquot, most of the protein N-termini identified across the two single-step elution conditions (**Figure 2B**). This result demonstrates that sequential two-step elution retains the highly selective Fraction 1 while additionally recovering strongly retained N-terminal peptides in Fraction 2.

**Figure 2.**
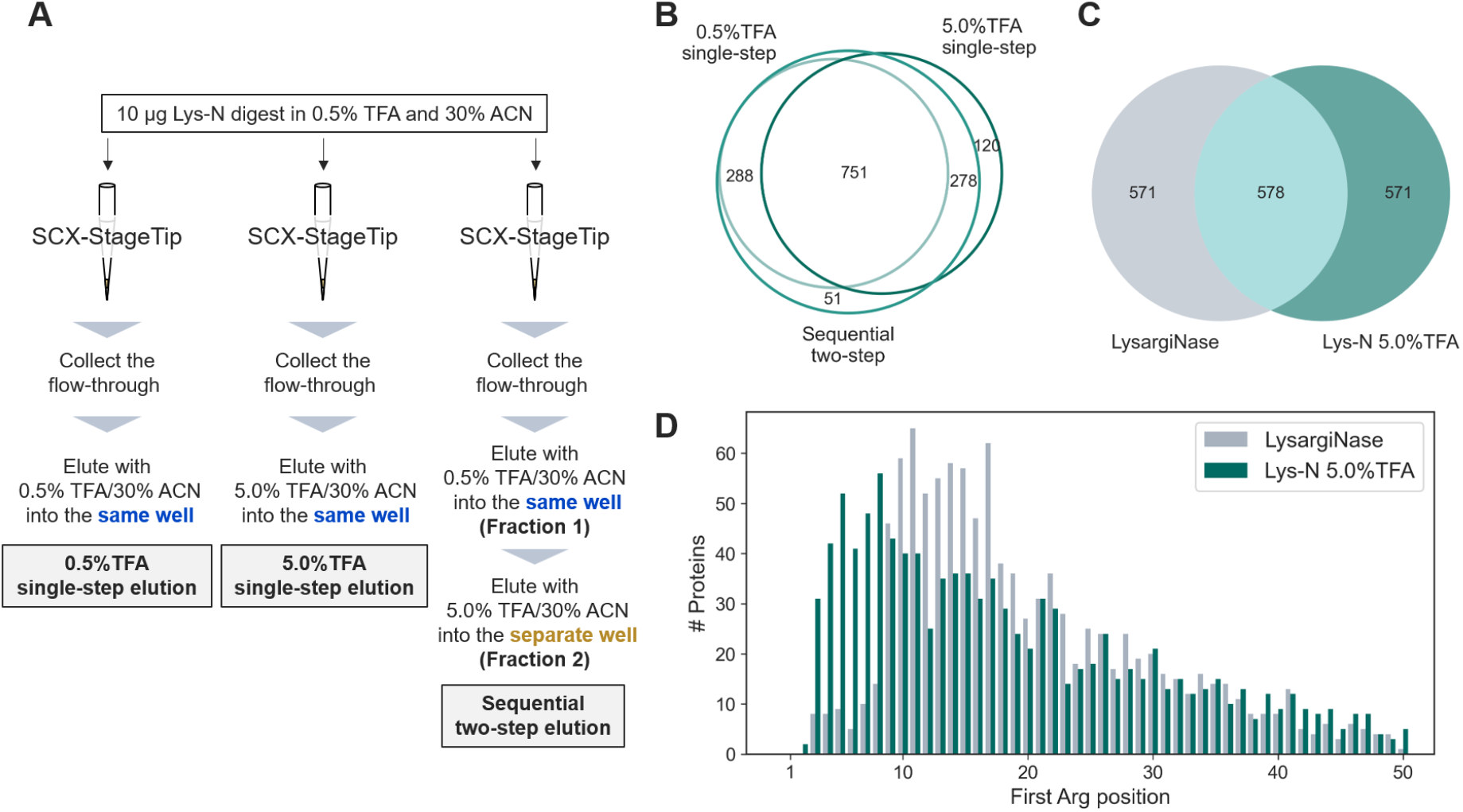
Comparison of SCX elution formats for LysN-based CHAMP-N and evaluation of complementarity with conventional LysargiNase-based CHAMP-N. (A) Schematic comparison of single-step elution and sequential two-step SCX elution. All samples were loaded in 30% ACN containing 0.5% TFA. For the two single-step elution conditions, separate aliquots of the same LysN digest were loaded onto separate SCX-StageTips. One StageTip was eluted with 0.5% TFA in 30% ACN, whereas the other was eluted with 5.0% TFA in 30% ACN. For each condition, the flow-through and corresponding eluate were combined into a single fraction, and each combined fraction was analyzed separately. For sequential two-step elution, the flow-through and 0.5% TFA eluate from a single StageTip were collected as Fraction 1. The same StageTip was then eluted with 5.0% TFA in 30% ACN into a separate well to obtain Fraction 2. The two fractions were analyzed separately, and their identification results were combined. (B) Overlap between protein N-termini identified in the union of the two independently analyzed single-step elution conditions and those identified after combining the identification results from the two sequential fractions. (C) Overlap between protein N-termini identified using conventional LysargiNase-based CHAMP-N and those identified using LysN-based CHAMP-N with 5.0% TFA single-step elution. (D) Distribution of first Arg positions in the protein sequences corresponding to protein N-termini identified using conventional LysargiNase-based CHAMP-N or LysN-based CHAMP-N with 5.0% TFA single-step elution. The first Arg position was defined as the residue number of the most N-terminal Arg in each protein sequence. For panels B-D, only N-terminal peptides identified in at least two of three technical replicates were included. Each leading razor protein associated with at least one identified protein N-terminal peptide was counted as one protein N-terminus.

These results revealed a trade-off between expanded coverage of protein N-termini and N-terminal peptide selectivity as the TFA concentration increased. Single-step elution with 0.5% TFA provided high selectivity, whereas single-step elution with 5.0% TFA increased the coelution of internal peptides but also recovered strongly retained N-terminal peptides, yielding the largest number of identified protein N-termini among the single-step conditions. Sequential two-step elution separated these peptide populations into two fractions from a single sample aliquot and increased cumulative coverage when the identification results were combined. However, this approach required two nanoLC/MS/MS runs. For the subsequent comparison with conventional LysargiNase-based CHAMP-N, we therefore used single-step elution with 5.0% TFA, which provided the largest number of identified protein N-termini in a single run, allowing the workflows to be compared using the same number of nanoLC/MS/MS measurements.

Using this condition, we compared LysN-based CHAMP-N with conventional LysargiNase-based CHAMP-N. LysN-based CHAMP-N identified 1,149 protein N-termini, whereas conventional LysargiNase-based CHAMP-N identified 1,149 protein N-termini (**Figure 2C**). Of these, 578 protein N-termini were identified under both conditions, whereas 571 and 571 protein N-termini were unique to the conventional LysargiNase and LysN conditions, respectively. Combining the identifications from both conditions yielded 1,720 protein N-termini, representing a 49.7% increase over conventional LysargiNase-based CHAMP-N alone. The partial overlap demonstrates that LysN-based CHAMP-N accesses a complementary set of protein N-termini rather than merely reproducing those detected by conventional LysargiNase-based CHAMP-N.

To examine the relationship between this complementarity and Arg residues near the protein N-terminus, we defined the first Arg position as the residue number of the most N-terminal Arg in each protein sequence and compared its distribution between the two conditions (**Figure 2D**). Protein N-termini identified under the LysN condition were more frequently derived from proteins with first Arg positions close to the protein N-terminus than those identified under the conventional LysargiNase condition. This result is consistent with the design principle that cleavage at a proximal Arg by LysargiNase generates a short N-terminal peptide, whereas LysN bypasses Arg and extends the peptide to the first downstream Lys. For example, the N-terminal peptides ADPRVRQI from tubulin-specific chaperone A (UniProt accession O75347) and MQRASRL from ubiquitin-conjugating enzyme E2 T (UniProt accession Q9NPD8) were identified exclusively under the LysN condition. Conventional LysargiNase digestion would be expected to generate the three-residue peptide ADP and the two-residue peptide MQ, respectively, which are too short for reliable LC/MS/MS identification. These examples illustrate how LysN-based CHAMP-N recovers protein N-termini that are poorly accessible under conventional LysargiNase digestion, thereby providing complementary coverage while retaining a single-step SCX workflow.

### ArgN-like CHAMP-N expands coverage of protein N-termini by suppressing Lys-directed cleavage

Next, to avoid cleavage at Lys residues near the protein N-terminus while retaining cleavage N-terminal to Arg, we performed ArgN-like CHAMP-N, in which proteins were D3-acetylated with acetic anhydride-d6 before LysargiNase digestion and SCX separation (**Figure 3A**). The principle of suppressing Lys-dependent cleavage by LysargiNase through propionylation of Lys to obtain ArgN-like digestion has previously been reported in global proteomics.^39^ Stable-isotope chemical acetylation of free protein N-termini has also been used to quantify N-terminal acetylation stoichiometry.^40^ In this study, we applied D3-acetylation-mediated ArgN-like digestion to CHAMP-N and investigated whether N-terminal peptides could be extended beyond proximal Lys residues while retaining the N-terminal Arg residues of internal peptides required for SCX separation.

**Figure 3.**
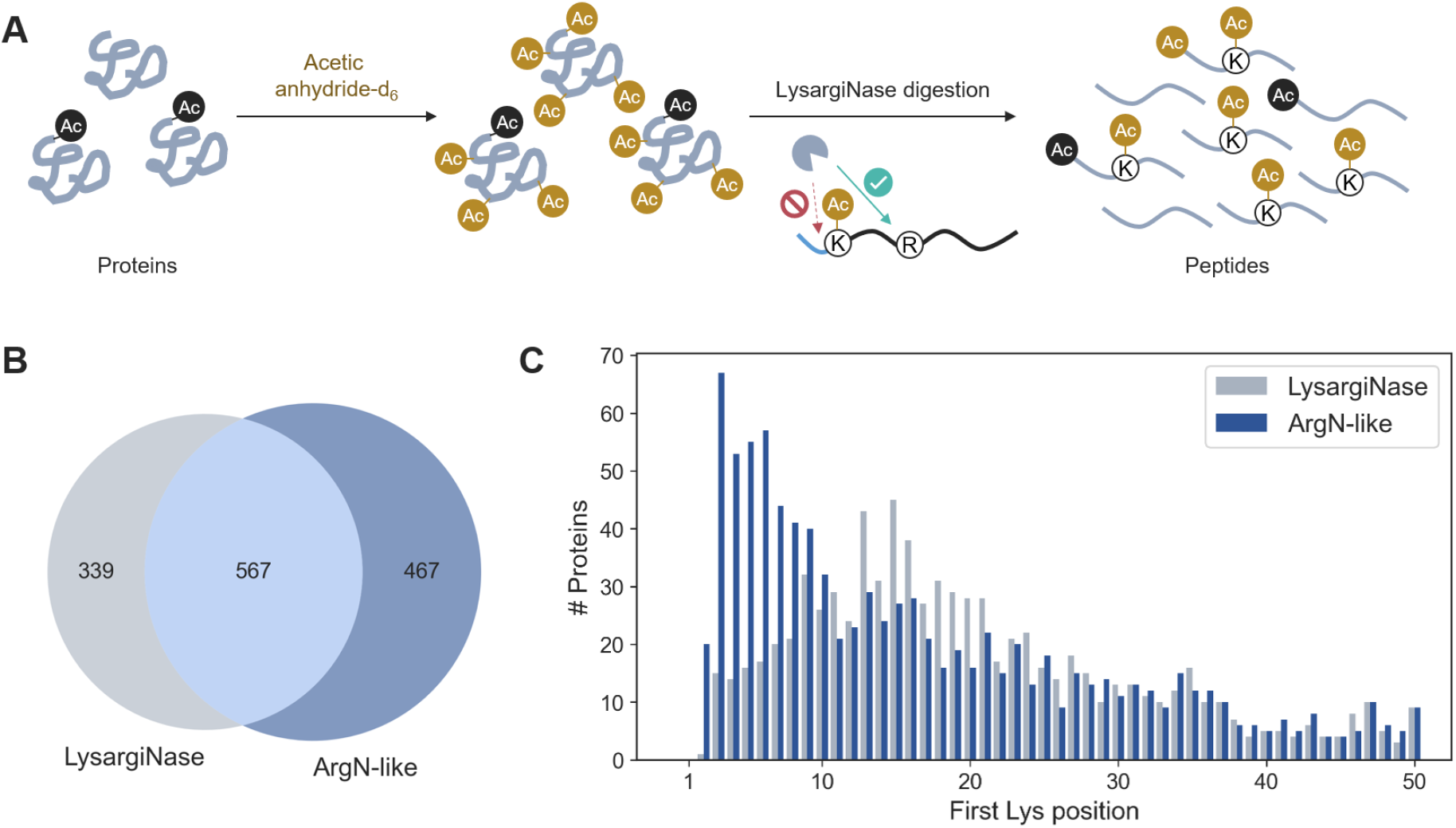
ArgN-like CHAMP-N expands coverage of protein N-termini by suppressing Lys-directed cleavage. (A) Schematic workflow of ArgN-like CHAMP-N, comprising D3-acetylation followed by LysargiNase digestion and SCX separation. Free protein N-termini and Lys side-chain ε-amino groups were derivatized with acetic anhydride-d6. Endogenous Nα-acetylation is shown as light acetylation (black Ac), whereas chemically introduced D3-acetylation is shown as heavy acetylation (gold Ac). Acetylation of Lys side chains suppresses LysargiNase cleavage at Lys residues, extending N-terminal peptides to the residue preceding the downstream Arg. (B) Overlap of protein N-termini identified under the conventional LysargiNase and ArgN-like conditions. (C) Distribution of first Lys positions in the protein sequences corresponding to protein N-termini identified under the conventional LysargiNase and ArgN-like conditions. The first Lys position was defined as the position of the first Lys residue from the protein N-terminus. Bars indicate the number of protein N-termini at each position. For panels B and C, only N-terminal peptides identified in at least two of three technical replicates were included, and each leading razor protein associated with at least one identified protein N-terminal peptide was counted as one protein N-terminus.

This treatment introduced heavy acetyl groups onto the α-amino groups of initially free protein N-termini and the ε-amino groups of Lys side chains, whereas endogenously Nα-acetylated N-termini retained their light acetyl groups. Acetylation of Lys side chains suppressed cleavage N-terminal to Lys, allowing N-terminal peptides to extend beyond proximal Lys residues to the residue immediately preceding the first downstream Arg. Cleavage at Arg generated internal peptides beginning with Arg, thereby preserving the charge characteristics required for SCX separation in CHAMP-N.

Under the conventional LysargiNase condition, 906 protein N-termini were identified, whereas 1,034 protein N-termini were identified under the ArgN-like condition (**Figure 3B**). Of these, 567 protein N-termini were identified under both conditions, whereas 339 and 467 protein N-termini were uniquely identified under the conventional LysargiNase and ArgN-like conditions, respectively. The ArgN-like condition alone yielded 14.1% more protein N-termini than the conventional LysargiNase condition. Combining the identifications from both conditions yielded 1,373 protein N-termini, representing a 51.5% increase in cumulative coverage of protein N-termini over the conventional LysargiNase condition alone. These results demonstrate that suppression of Lys-directed cleavage by LysargiNase generates protein N-terminus identifications complementary to those obtained under the conventional LysargiNase condition.

To examine the relationship between this complementarity and Lys residues near the protein N-terminus, we defined the first Lys position as the residue number of the most N-terminal Lys in each protein sequence and compared its distribution between the two conditions (**Figure 3C**). Protein N-termini identified under the ArgN-like condition were more frequently derived from proteins with first Lys positions close to the protein N-terminus than those identified under the conventional LysargiNase condition, with the increase being particularly pronounced when the first Lys occurred within the first 10 residues. This result is consistent with D3-acetylation suppressing cleavage at proximal Lys residues and allowing N-terminal peptides to extend to the first downstream Arg. For example, the N-terminal peptides GKKGKVGKS from pre-rRNA 2′-O-ribose RNA methyltransferase FTSJ3 (UniProt accession Q8IY81) and PSKKKKYNA from Dr1-associated corepressor (UniProt accession Q14919) were identified exclusively under the ArgN-like condition. Conventional LysargiNase digestion would be expected to generate the single-residue peptide G and the two-residue peptide PS, respectively, which are too short for reliable LC/MS/MS identification. These examples illustrate how ArgN-like CHAMP-N recovers protein N-termini that are poorly accessible under conventional LysargiNase digestion by preventing cleavage at proximal Lys residues.

Acetylation of free N-termini and Lys side chains neutralizes protonatable amino groups and may therefore affect peptide precursor charge states. We compared the precursor charge-state distributions of N-terminal peptides identified under the conventional LysargiNase and ArgN-like conditions (**Figure S3**). Doubly charged precursor ions predominated under both conditions, and no clear shift toward lower charge states was observed under the ArgN-like condition. Thus, at least among the identified N-terminal peptides, D3-acetylation did not substantially alter the precursor charge-state distribution.

We next evaluated the residue selectivity of D3-acetylation. An exploratory open search using FragPipe first revealed +45.0294 Da mass shifts potentially localized to several residues other than the intended targets.^36,37^ Based on these observations, we performed a targeted variable-modification search in which the +45.0294 Da mass shift was allowed on Ser, Thr, Tyr, Arg, and His in addition to protein N-termini and Lys residues. When the same modified site in the same protein was identified in multiple peptides, it was counted only once. Among all unique sites assigned the +45.0294 Da mass shift, 15.23% and 71.78% were assigned to protein N-termini and Lys residues, respectively. Assignments to Ser, Thr, Tyr, His, and Arg accounted for 8.17%, 3.34%, 0.73%, 0.68%, and 0.07%, respectively. Thus, 87.01% of the assignments occurred at the intended targets, whereas 12.99% occurred at other residues, indicating that chemical D3-acetylation was not completely selective. Although these additional modifications can complicate acetylation-site localization in some peptide-spectrum matches, protein N-terminal peptides were classified according to their sequences and positions within the corresponding proteins. Therefore, localization uncertainty does not invalidate the N-terminal sequence assignments used for the coverage analysis.

The apparent D3-acetylation rates of annotated protein N-termini that were free before chemical treatment and of Lys residues were 90.3% and 92.8%, respectively. These high modification rates are consistent with efficient derivatization of the intended sites and effective suppression of Lys-directed cleavage by LysargiNase. Nevertheless, the targeted search produced apparent off-target assignments, particularly at Ser and Thr, and complete residue selectivity could not be established. Excluding these candidate modifications from database searches may result in missed identifications, whereas including them as variable modifications increases the search space. Residue selectivity and modification localization therefore remain limitations of the present chemical acetylation-based approach.

### Complementary cleavage control expands coverage of protein N-termini

Finally, we directly compared conventional LysargiNase-based CHAMP-N, LysN-based CHAMP-N using 5.0% TFA single-step elution, and ArgN-like CHAMP-N and evaluated the cumulative coverage of protein N-termini obtained by integrating identifications from the three workflows (**Figure 4**).

**Figure 4.**
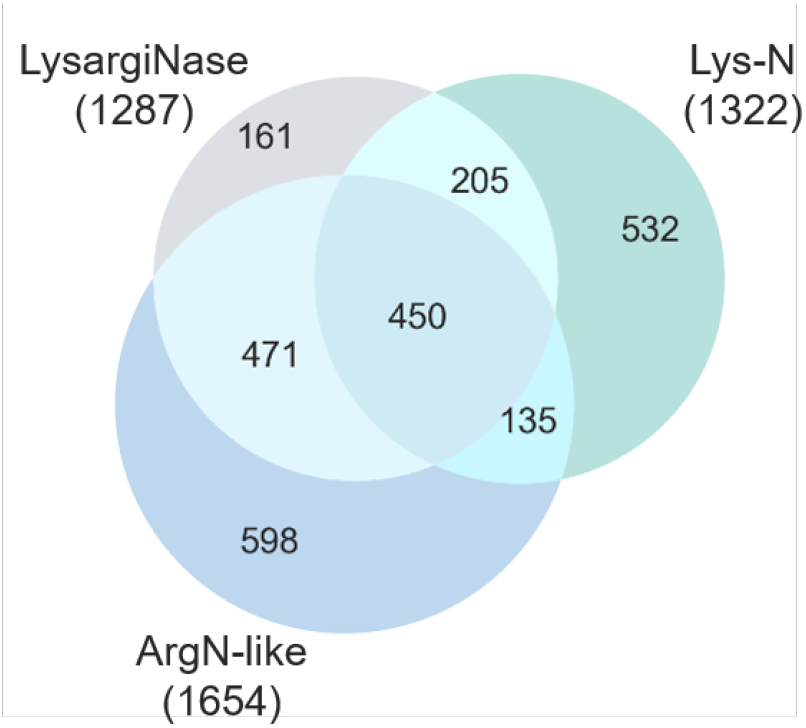
Integration of complementary CHAMP-N workflows expands coverage of protein N-termini. Overlap among protein N-termini identified using conventional LysargiNase-based CHAMP-N, LysN-based CHAMP-N with 5.0% TFA single-step SCX elution, and ArgN-like CHAMP-N (D3-acetylation followed by LysargiNase digestion). Numbers in parentheses indicate the total number of protein N-termini identified by each workflow. Only N-terminal peptides identified in at least two of three technical replicates were included. Each leading razor protein associated with at least one identified protein N-terminal peptide was counted as one protein N-terminus. All samples were analyzed using the same extended LC gradient and an MS2 scan of the first mass at *m*/*z* 100, as described in the Materials and Methods.

The numbers of identified protein N-termini were 1,287, 1,322, and 1,654 under the conventional LysargiNase, LysN, and ArgN-like conditions, respectively. Of these, 450 protein N-termini were common to all three conditions, whereas 161, 532, and 598 protein N-termini were unique to the conventional LysargiNase, LysN, and ArgN-like conditions, respectively. Combining the identifications from all three conditions yielded 2,552 protein N-termini, adding 1,265 protein N-termini to the 1,287 identified under the conventional LysargiNase condition alone. Thus, cumulative coverage of protein N-termini was doubled relative to the conventional condition.

LysN digestion bypasses cleavage at Arg residues near the protein N-terminus, extending N-terminal peptides to the residue immediately preceding the first downstream Lys. By contrast, in ArgN-like digestion, D3-acetylation suppresses Lys-directed cleavage by LysargiNase, extending N-terminal peptides to the residue immediately preceding the first downstream Arg. Accordingly, both digestion conditions generate N-terminal peptides with C-terminal boundaries different from those generated by conventional LysargiNase digestion. The large numbers of protein N-termini uniquely identified under the LysN and ArgN-like conditions indicate that selectively using Lys- or Arg-directed cleavage can reduce the sequence-dependent detection bias associated with any single digestion condition.

Quantitative reproducibility was assessed for each workflow by calculating the relative standard deviations (RSDs) of MS intensities for N-terminal peptides identified in all three technical replicates (**Figure S4**). The median RSD was approximately 15% under each condition, and the distributions were broadly similar. Thus, neither LysN-based CHAMP-N using 5.0% TFA single-step elution nor ArgN-like CHAMP-N compromised quantitative reproducibility relative to conventional LysargiNase-based CHAMP-N.

Because LysN and ArgN-like digestion can preserve Arg and Lys residues, respectively, within protein N-terminal peptides, these complementary cleavage conditions may extend the applicability of Lys/Arg SILAC to N-terminal peptide quantification. For each identified protein N-terminus, we defined sequence-level compatibility with Lys/Arg SILAC as the presence of at least one Lys or Arg in at least one assigned annotated protein N-terminal peptide. The proportion of protein N-termini meeting this criterion was 49.3% under the conventional LysargiNase condition (635/1,287 protein N-termini), 74.7% under the LysN condition (987/1,322 protein N-termini), 68.9% under the ArgN-like condition (1,140/1,654 protein N-termini), and 75.3% when identifications from all three conditions were combined (1,922/2,552 protein N-termini). However, this assessment reflects only the sequences of the identified peptides; labeling efficiency and quantitative performance in SILAC samples were not experimentally evaluated.

One limitation of ArgN-like CHAMP-N is that D3-acetylation requires an additional chemical reaction and can generate off-target modifications. If a protease that selectively cleaves N-terminal to Arg becomes available, ArgN-like cleavage specificity could potentially be achieved without chemical derivatization. Another limitation is that the present strategies are designed primarily to rescue N-terminal peptides that are too short and do not directly address peptides that remain excessively long after digestion. However, in silico digestion predicted that only 2.05% of the N-terminal peptides generated by conventional LysargiNase digestion would be 50 residues or longer. Thus, at least under the conventional digestion condition, the population potentially rescued by secondary proteolysis after N-terminal peptide separation is expected to be small, suggesting that the resulting gain in coverage would be limited.

Overall, complementary cleavage control using LysN and ArgN-like digestion doubled coverage of protein N-termini without compromising quantitative reproducibility and increased the proportion of identified protein N-termini potentially amenable to Lys/Arg SILAC.

## Conclusions

This study addresses a fundamental sequence-dependent limitation of N-terminomics: cleavage near the protein N-terminus generates peptides too short for LC/MS/MS analysis. LysN and ArgN-like digestion shifted the C-terminal boundaries of these peptides while preserving the N-terminal basic residues of internal peptides required for SCX enrichment. Combining the three workflows increased cumulative coverage from 1,287 to 2,552 protein N-termini, nearly twofold, without compromising technical reproducibility. These results establish coordinated control of cleavage specificity and SCX selectivity as a practical strategy for expanding the observable protein N-terminome. The current strategies mainly address losses caused by proximal Lys or Arg residues. ArgN-like CHAMP-N also relies on D3-acetylation, which was associated with apparent off-target modification assignments and increased ambiguity in modification localization and database searching. Development of a protease that cleaves selectively N-terminal to Arg would eliminate this chemical derivatization step and simplify the workflow. Additional enrichment-compatible cleavage conditions could extend the approach to other sequence contexts and facilitate more comprehensive analyses of alternative translation initiation, signal-peptide processing, and regulated proteolysis.

## Author Contributions

The conceptual design of the project was done by EK, KI, and YI. The experimental designs were conducted by AT and EK. Sample preparation, LC/MS/MS, and data analysis were performed by AT. The manuscript was written by AT, EK, and YI.

## Acknowledgments

We would like to thank members of the Department of Molecular Systems BioAnalysis in Kyoto University for fruitful discussions. This work was supported by AMED-PRIME (JP25gm7010003h0001) to EK, JST-ASTEP (JPMJTR25U9) to EK, Grants-in-Aid for Scientific Research (KAKENHI; Grants JP26K02156 and JP26K22909 to EK, JP25K22526, JP23H04924, and JP23K18185 to YI), Toyota Riken Scholarship to EK, and Takeda Science Foundation to EK.

## Supporting Information

The Supporting Information is available free of charge at https://XXXX. Supplementary methods; theoretical length distribution of N-terminal peptides generated by conventional LysargiNase digestion of canonical human protein sequences; optimization of TFA concentration for single-step SCX elution of N-terminal peptides from LysN digests; precursor charge-state distributions of N-terminal peptides under conventional LysargiNase and ArgN-like conditions; and quantitative reproducibility across complementary CHAMP-N workflows.

## Data availability statement

The MS raw data files and the analysis files containing the identification and quantification results for peptides and protein groups have been deposited at the ProteomeXchange Consortium (http://proteomecentral.proteomexchange.org) via the jPOST partner repository (https://jpostdb.org) with the dataset identifier JPST004881/PXD083406.^41,42^

## TOC

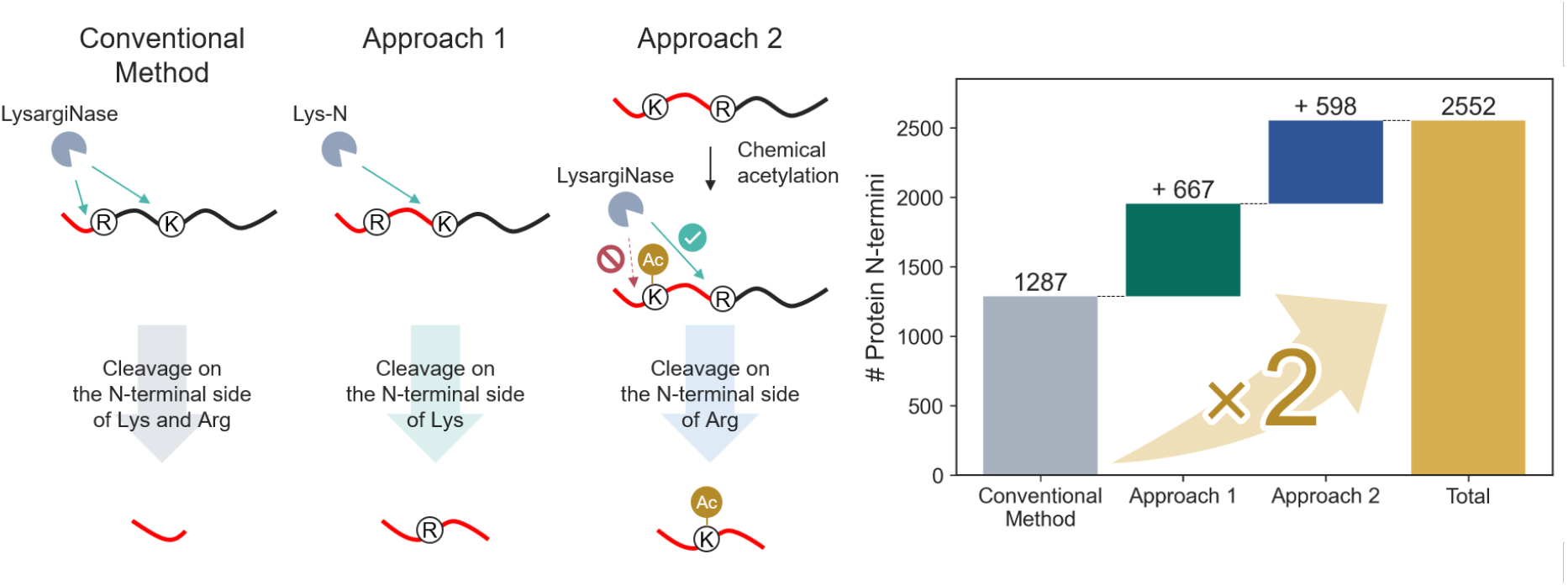

