## Supporting information for "Expanding N-terminomics Coverage by Extending N-terminal Peptide Length"

**Table of Contents**

**Supplementary Methods**: Open and targeted searches for D3-acetylation-compatible mass shifts

**Figure S1.** Theoretical length distribution of N-terminal peptides generated by conventional LysargiNase digestion of canonical human protein sequences.

**Figure S2.** Optimization of TFA concentration for single-step SCX elution of N-terminal peptides from Lys-N digests.

**Figure S3.** Precursor charge-state distributions of N-terminal peptides under conventionaLysargiNase and ArgN-like conditions.

**Figure S4.** Quantitative reproducibility across complementary CHAMP-N workflows.

**Supplementary Methods**

**Open and targeted searches for D3-acetylation-compatible mass shifts**

The raw data files from the ArgN-like CHAMP-N samples used for Figure 3 were analyzed using MSFragger (version 4.3) in FragPipe (version 23.1) and processed with Philosopher (version 5.1.2).[1,2] Spectra were searched against the same human UniProtKB/Swiss-Prot protein sequence database including isoforms as used for the MaxQuant analysis (release 2020.10; 42,373 sequences), supplemented with common contaminants and reversed decoy sequences. Enzyme specificity was set to "lysn_promisc" (cleavage N-terminal to Lys and Arg), with up to five missed cleavages. Carbamidomethylation of Cys was specified as a fixed modification. Acetylation of protein N-termini, D3-acetylation of protein N-termini (+45.0294 Da), D3-acetylation of Lys residues (+45.0294 Da), and oxidation of Met were specified as variable modifications. PSMs were filtered at a 1% FDR, and decoy and contaminant PSMs were excluded from subsequent analyses.

An exploratory open search was first performed to identify candidate mass shifts and their possible residue localization. The precursor-mass search range was −150 to +500 Da, and the fragment-ion mass tolerance was 20 ppm. PTM-Shepherd (version 3.0.2)[3] was used for mass-shift summarization and residue localization, with the "Localize mass shift" option enabled. Mass shifts within 0.01 Da of +45.0294 Da were classified as compatible with D3-acetylation.

Assignments to residues other than the intended protein N-termini and Lys residues were subsequently evaluated using a targeted variable-modification search. The precursor- and fragment-ion mass tolerances were ±20 and 20 ppm, respectively. In addition to D3-acetylation of protein N-termini and Lys residues, the +45.0294 Da mass shift was specified as a variable modification on Ser, Thr, Tyr, Arg, and His. All other search settings and FDR criteria were identical to those used for the open search.

For the site-level distribution analysis, repeated assignments of the +45.0294 Da mass shift to the same site in the same protein across multiple peptide identifications were counted only once. All resulting unique sites assigned the +45.0294 Da mass shift were used as the denominator. The proportions assigned to protein N-termini and Lys, Ser, Thr, Tyr, Arg, and His residues were calculated for each replicate, and the mean percentages across replicates were reported.

The apparent D3-acetylation rate at initially free annotated protein N-termini was calculated as the proportion of D3-acetylated forms among the identified D3-acetylated and unmodified annotated protein N-termini. The apparent D3-acetylation rate of Lys was calculated as the proportion of identified Lys residues assigned the +45.0294 Da mass shift among all identified Lys residues. For the coverage analysis, protein N-terminal peptides were classified according to their sequences and positions within the corresponding proteins, irrespective of the exact localization of the +45.0294 Da mass shift within the peptide.

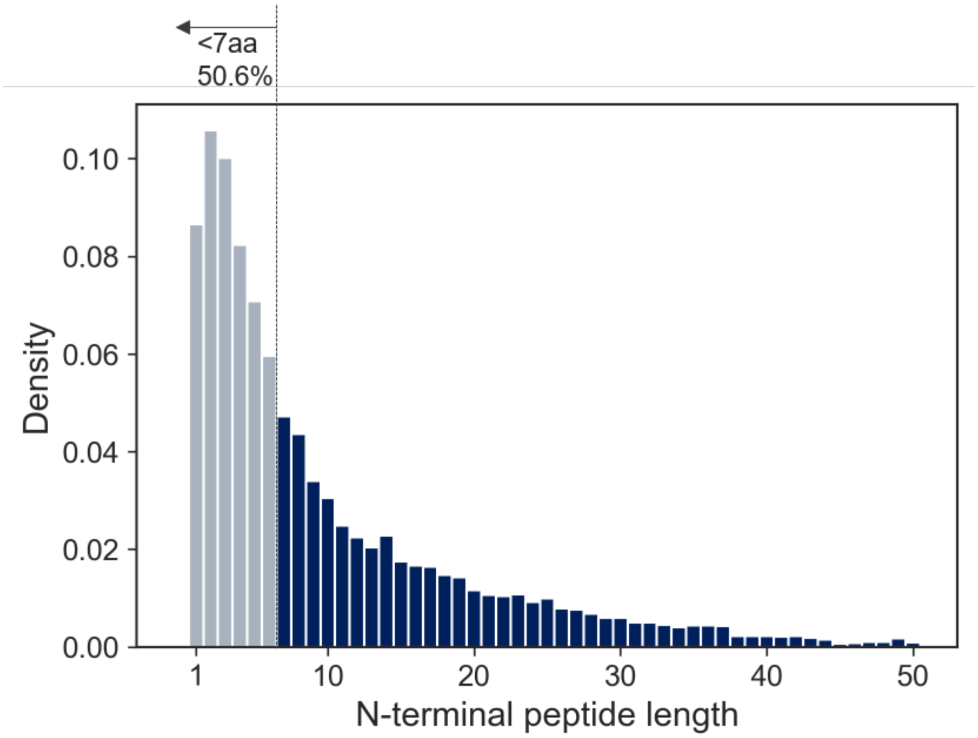


**Figure S1. Theoretical length distribution of N-terminal peptides generated by conventional LysargiNase digestion of canonical human protein sequences.** Canonical human protein sequences from UniProtKB/Swiss-Prot (release 2020.10, 20,385 proteins) were subjected to in silico conventional LysargiNase digestion, with cleavage modeled N-terminal to Lys and Arg. No missed cleavages were allowed, and initiator-methionine excision was not considered. The theoretical N-terminal peptide length was defined as the number of residues from the annotated protein N-terminus to the residue immediately preceding the first modeled cleavage site. Seven residues was used as the operational minimum because the experimental database searches used the default minimum peptide length of seven residues in MaxQuant (version 2.6.5.0). Predicted peptides shorter than seven residues, which were therefore outside the searchable peptide-length range, accounted for 50.6% of the theoretical N-terminal peptides and are shown in gray. Peptides of 50 residues or longer accounted for 2.05%. The seven-residue cutoff defines database-search eligibility, whereas the 50-residue cutoff is an operational length classification; neither represents an empirical lower or upper limit for LC/MS/MS detectability.

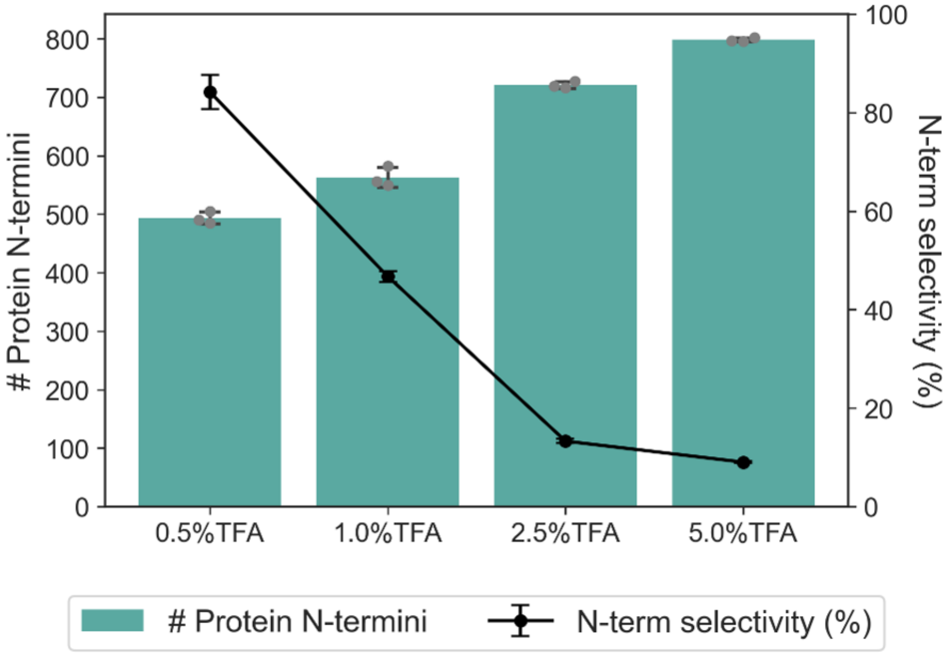


**Figure S2. Optimization of TFA concentration for single-step SCX elution of N-terminal peptides from Lys-N digests.** Aliquots, each equivalent to 10 µg of starting protein, were reconstituted in 30% ACN containing 0.5% TFA and loaded onto SCX StageTips. N-terminal peptides were eluted with 50 µL of 30% ACN containing 0.5%, 1.0%, 2.5%, or 5.0% TFA. For each condition, the flow-through and eluate were combined and analyzed by nanoLC/MS/MS. Bars indicate the mean number of identified protein N-termini. N-terminal peptide selectivity (%) was calculated as 100 × the summed MS intensity of N-terminal peptides divided by the summed MS intensity of all identified peptides. Gray dots represent individual technical replicates, and error bars indicate standard deviations (n = 3).

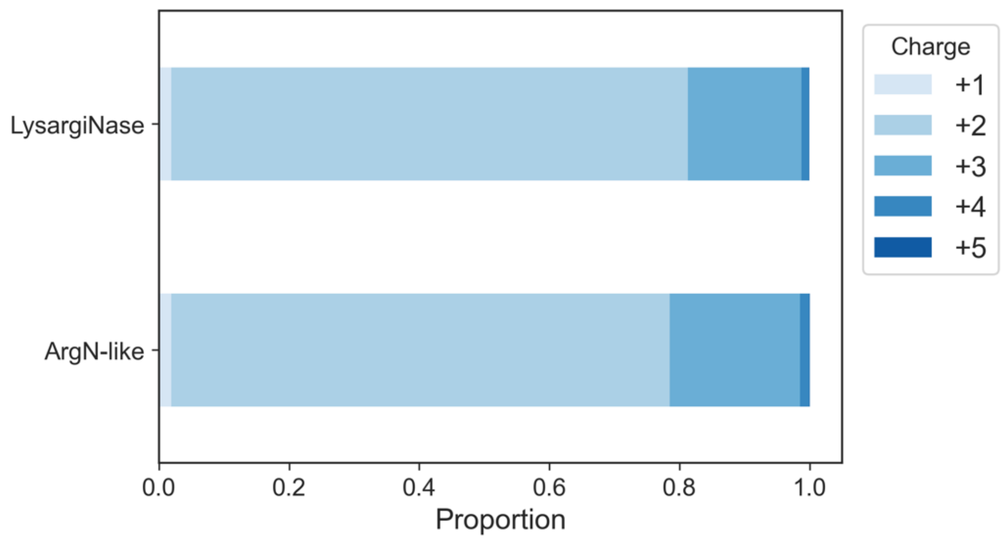


**Figure S3. Precursor charge-state distributions of N-terminal peptides under conventional LysargiNase and ArgN-like conditions.** Stacked bars show the relative proportions of precursor ions with charge states from +1 to +5 for identified N-terminal peptides. The conventional LysargiNase condition did not include chemical N-terminal acetylation, whereas the ArgN-like condition comprised D3-acetylation followed by LysargiNase digestion. Data from the three technical replicates used for Figure 3 were pooled within each condition.

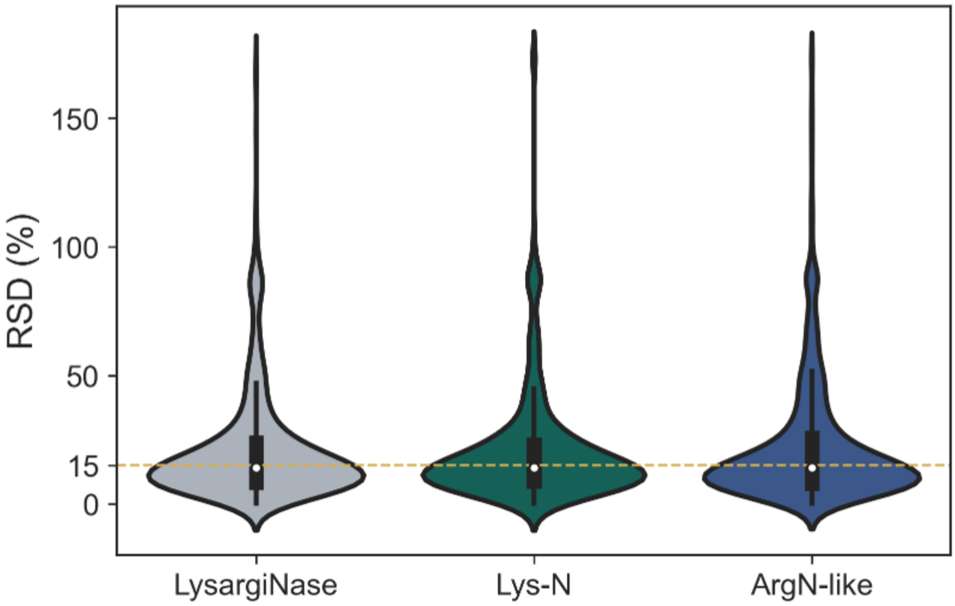


**Figure S4. Quantitative reproducibility across complementary CHAMP-N workflows.** Violin plots show the distributions of relative standard deviations (RSDs) for the MS intensities of N-terminal peptides obtained using conventional LysargiNase-based CHAMP-N, Lys-N-based CHAMP-N with 5.0% TFA single-step SCX elution, and ArgN-like CHAMP-N (D3-acetylation followed by LysargiNase digestion). Only N-terminal peptides identified in all three technical replicates used for Figure 4 were included. For each peptide, RSD (%) was calculated as 100 × the standard deviation divided by the mean of its MS intensities across the three technical replicates. The dashed line indicates an RSD of 15%.
